## Supplementary Figures for "A tuneable minimal cell membrane reveals that two lipid species suffice for life"

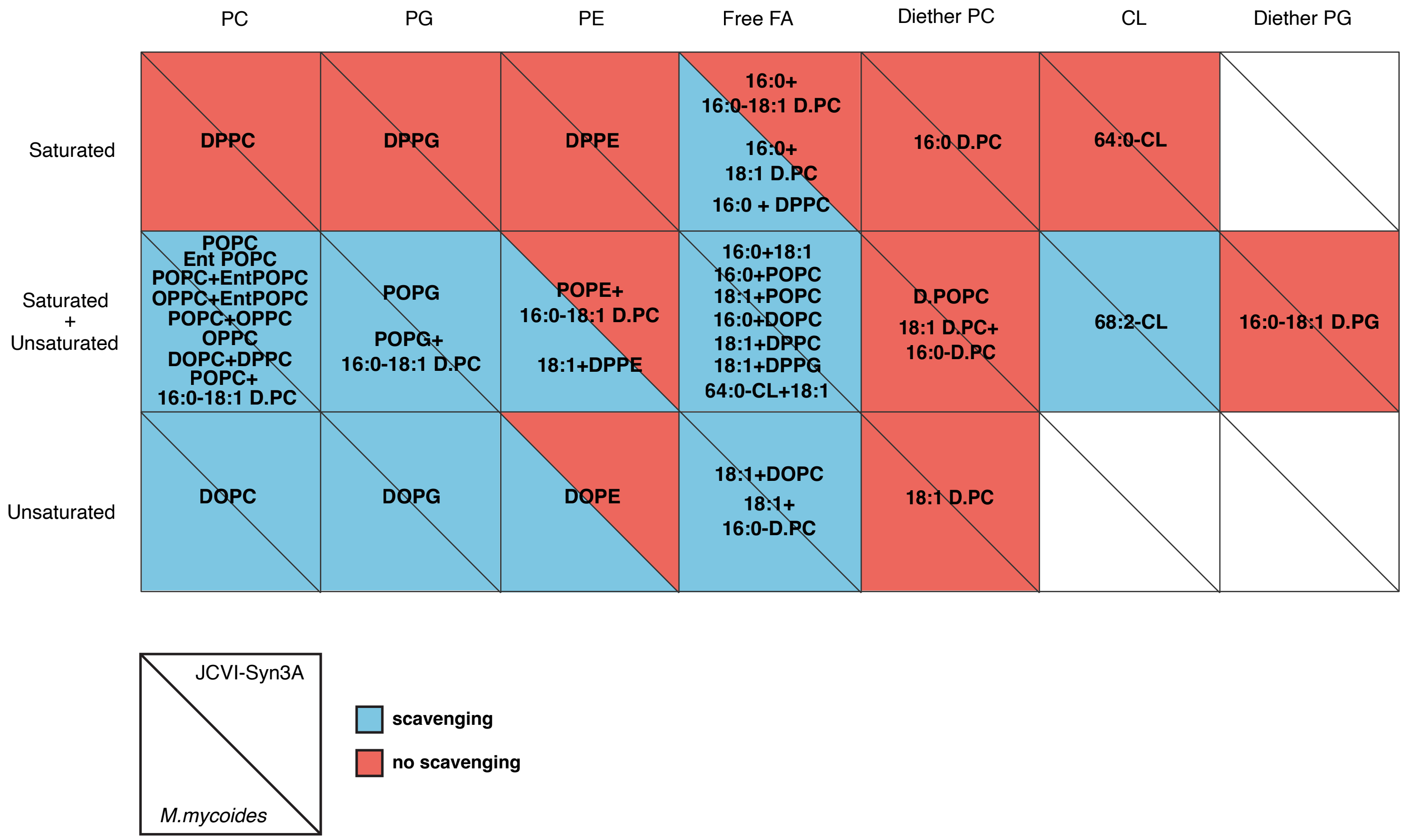


**Figure S1: Certain Lipid Features Determine Cells' Ability to Scavenge Acyl Chains**

All of the lipid diets fed to *M. mycoides* and JCVI-Syn3A were sorted based on lipid features, specifically by head group and acyl chain saturation, and the ability of *M. mycoides* and JCVI-Syn3A to scavenge acyl chains was determined by TLC analysis (Figure S4), with production of PG and Cardiolipin being taken as proof of scavenging occurring. As a result of this sorting, certain lipid features that lead to scavenging capacity, or the lack thereof, could be looked at. When only saturated phospholipids are fed, neither *M. mycoides* nor JCVI-Syn3A can scavenge acyl chains. When free 16:0 is added to the diet *M. mycoides* is able to produce PG and Cardiolipin, while JCVI-Syn3A is not. Furthermore, JCVI-Syn3A is unable to scavenge acyl chains from PE species of lipid, regardless of saturation.

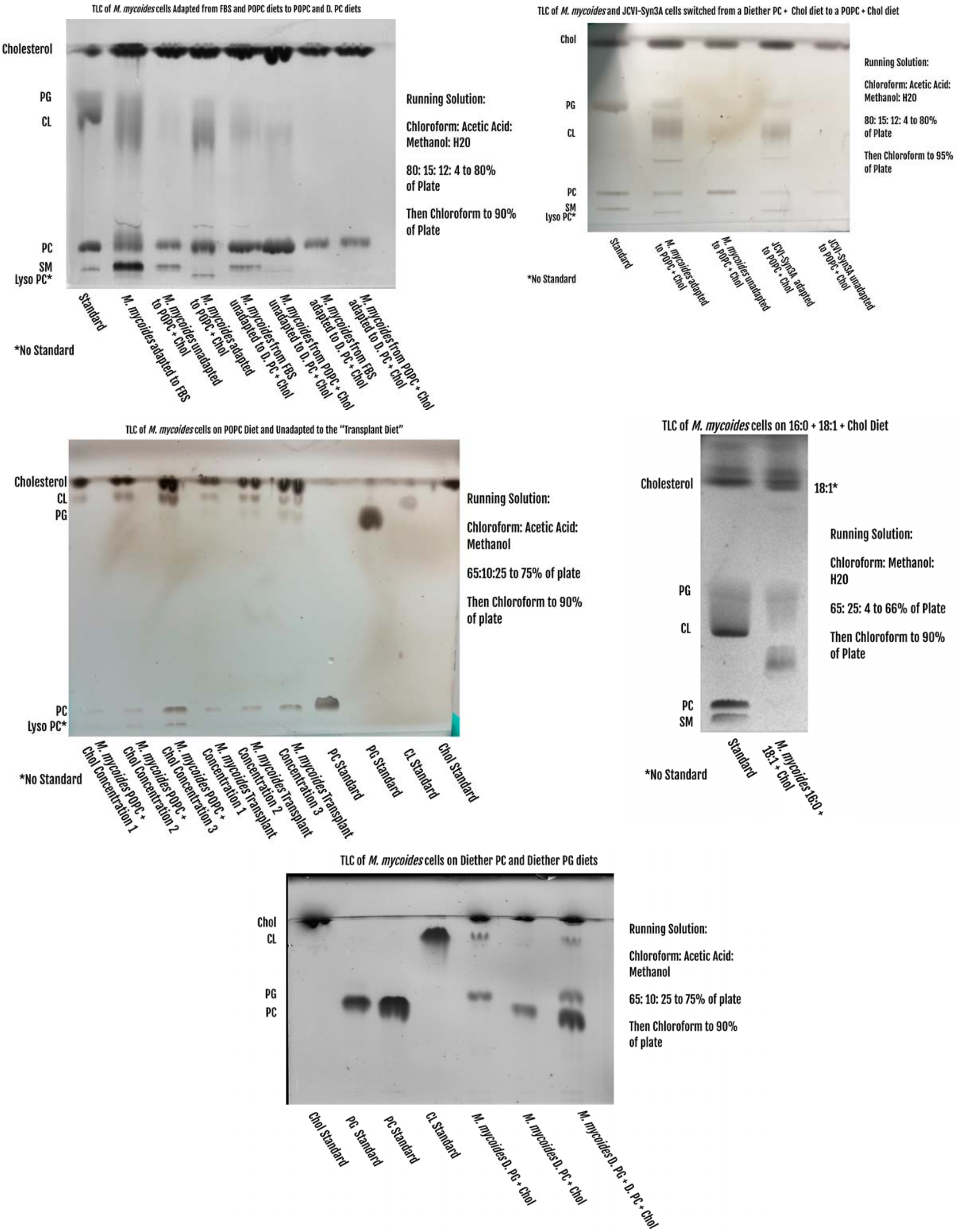


**Figure S2: TLC Analysis of Lipid Classes Present in Whole Cell Lipid Extracts**

The synthesis of PG and Cardiolipin by *M. mycoides* and JCVI-Syn3A fed on diets lacking free fatty acids provides evidence of acyl chain scavenging, as fatty acids are required to synthesize these lipid classes. To assay PG and Cardiolipin production, whole cell lipid extractions were performed on both organisms, which were then separated and visualized using thin layer chromatography against lipid standards to determine the presence or absence of PG and Cardiolipin. All plates are annotated.

**S3a**

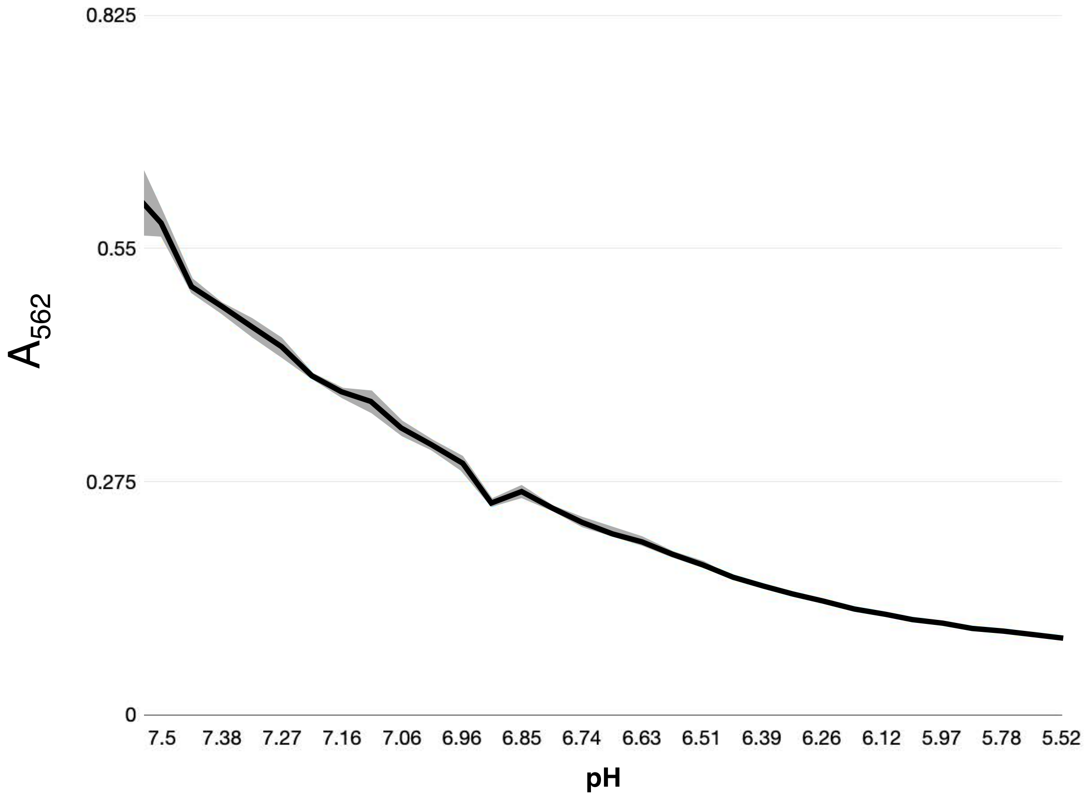


**S3b**

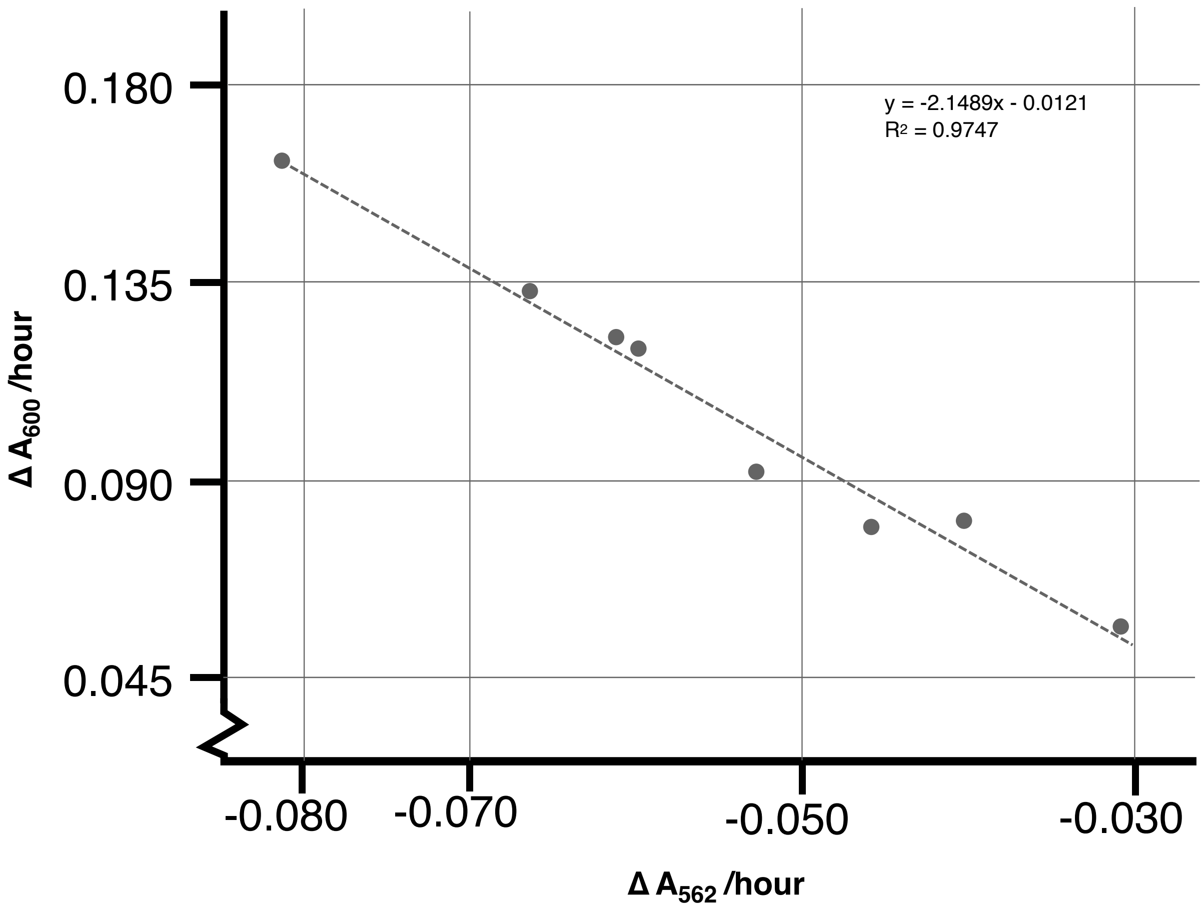


**Figure S3: Validating Phenol Red Absorbance Rates as a Method to Measure Cell Growth**

**a** The absorbance of phenol red in SP4 media at 562nm ($A_{562}$) as a function pH measured with an acid titration shows how phenol red signal $A_{562}$ decreases as media is acidified. **b** A scatterplot showing the relationship between growth rate values for *M. mycoides* obtained by measuring the optical density of cells at 600nm ($OD_{600}$) and metabolic growth index rates obtained by measuring changes in pH by looking at phenol red $A_{562}$. Eight diets were chosen to span a range of lipid species, saturations, and inclusions or exclusions of free fatty acids to ensure that they were representative of all diets. Measuring growth with phenol red $A_{562}$ was chosen due to difficulties obtaining reproducible $OD_{600}$ growth rates for JCVI-Syn3A. Furthermore, differences in cell size and growth behavior lead to difficulties properly comparing *M. mycoides* and JCVI-Syn3A values obtained from $OD_{600}$. Growth rates obtained by $OD_{600}$ are consistently ~2-fold higher than those obtained from phenol red $A_{562}$.

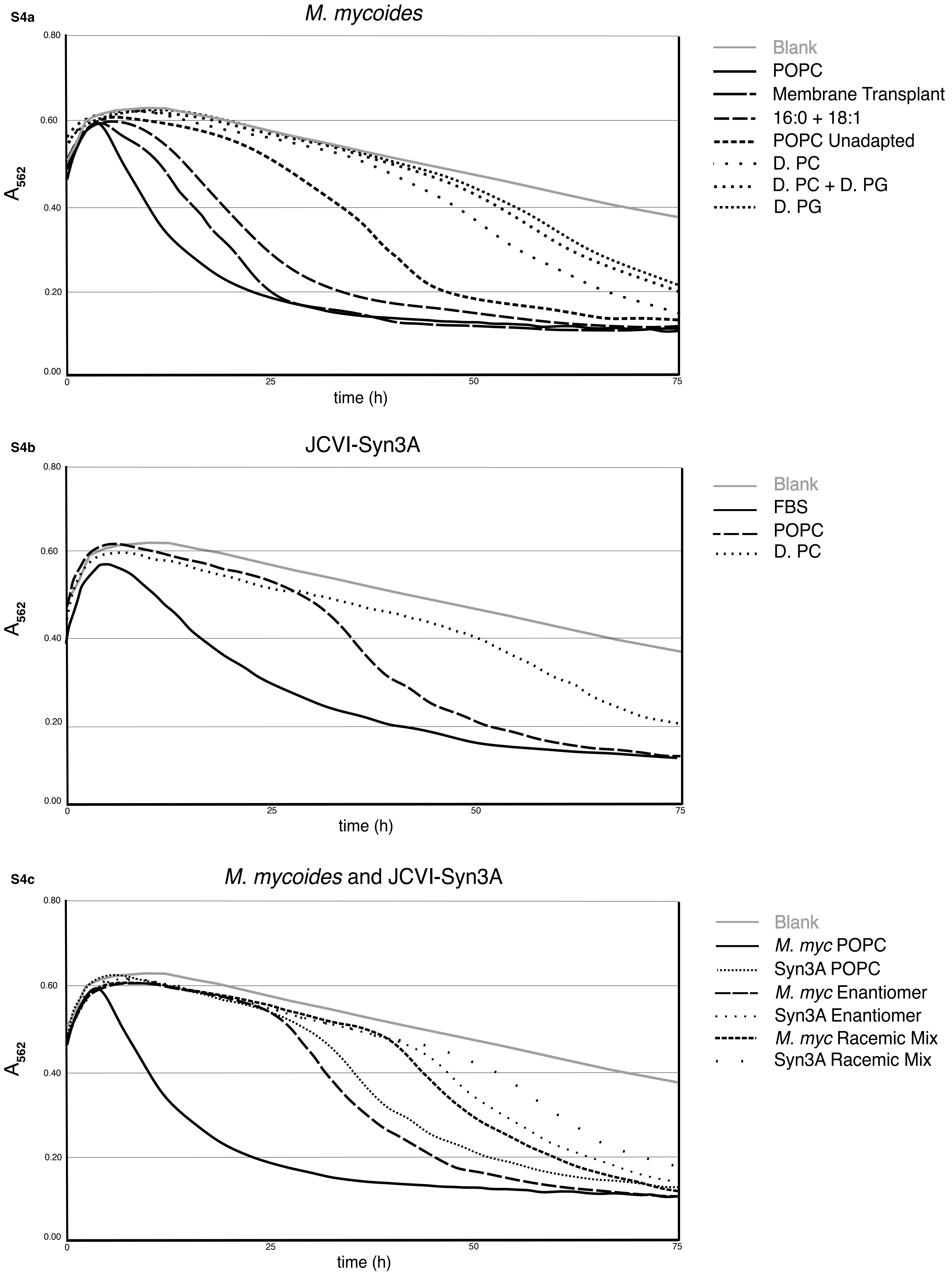


**Figure S4: Full $A_{562}$ Growth Curves**

A plot of the full average $A_{562}$ growth curves for all conditions in Figures 3, 4, and 5. Growth rates were calculated by taking an exponential fit of the most exponential part of each growth curve.

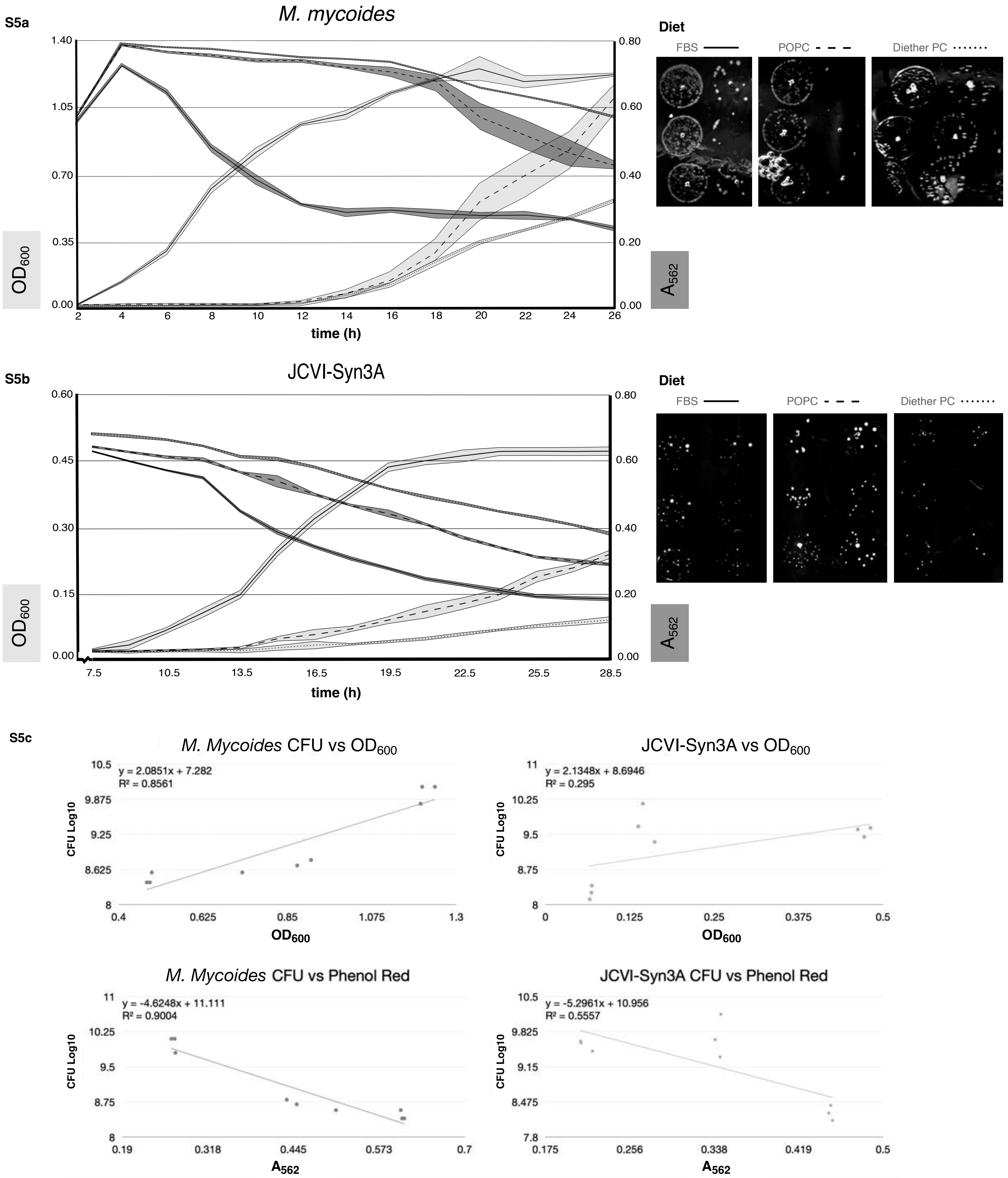


**Figure S5: Growth of *M. mycoides* and JCVI-Syn3A manually measured by $OD_{600}$, phenol red $A_{562}$, and Colony-Forming Units**

**a** Growth curves for *M. mycoides* on FBS, POPC, and Diether PC diets in phenol red media obtained by manual measurements of $OD_{600}$ and $A_{562}$. At 24 hours, cells were taken for each condition and serially diluted on a SP4+FBS plate to obtain a count of colony-forming units. **b** Growth curves for JCVI-Syn3A cells on FBS, POPC, and Diether PC diets in phenol red media obtained by manual measurements of $OD_{600}$ and $A_{562}$. At 24 hours, cells were taken for each condition and serially diluted on a SP4+FBS plate to obtain a count of colony-forming units. **c** Comparisons between CFUs, $OD_{600}$ and $A_{562}$ readings show a correlation between higher $OD_{600}$ and $A_{562}$ values and more CFUs, validating the absorbance-based methods of measuring growth. The correlation was weakest for JCVI-Syn3A cells measured by $OD_{600}$, further validating the decision to use phenol red $A_{562}$-based growth assays.

S6a

**FBS overview**

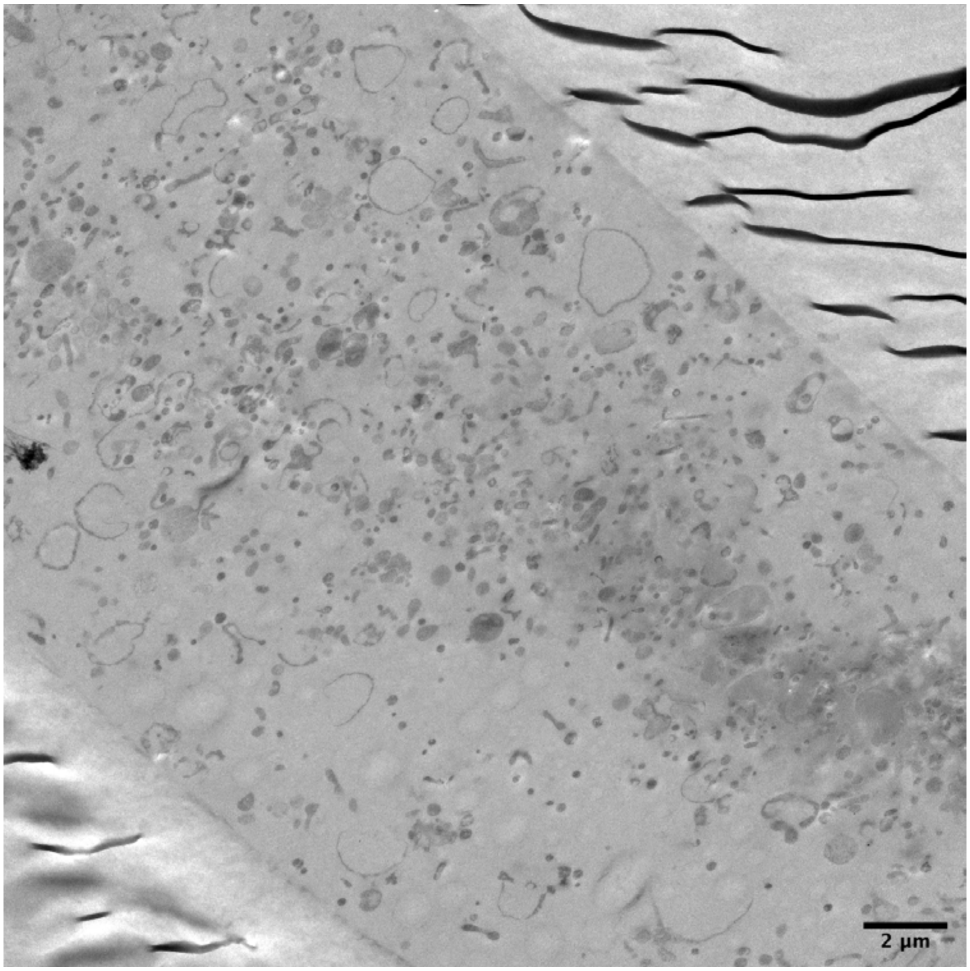


**FBS overview with cell count**

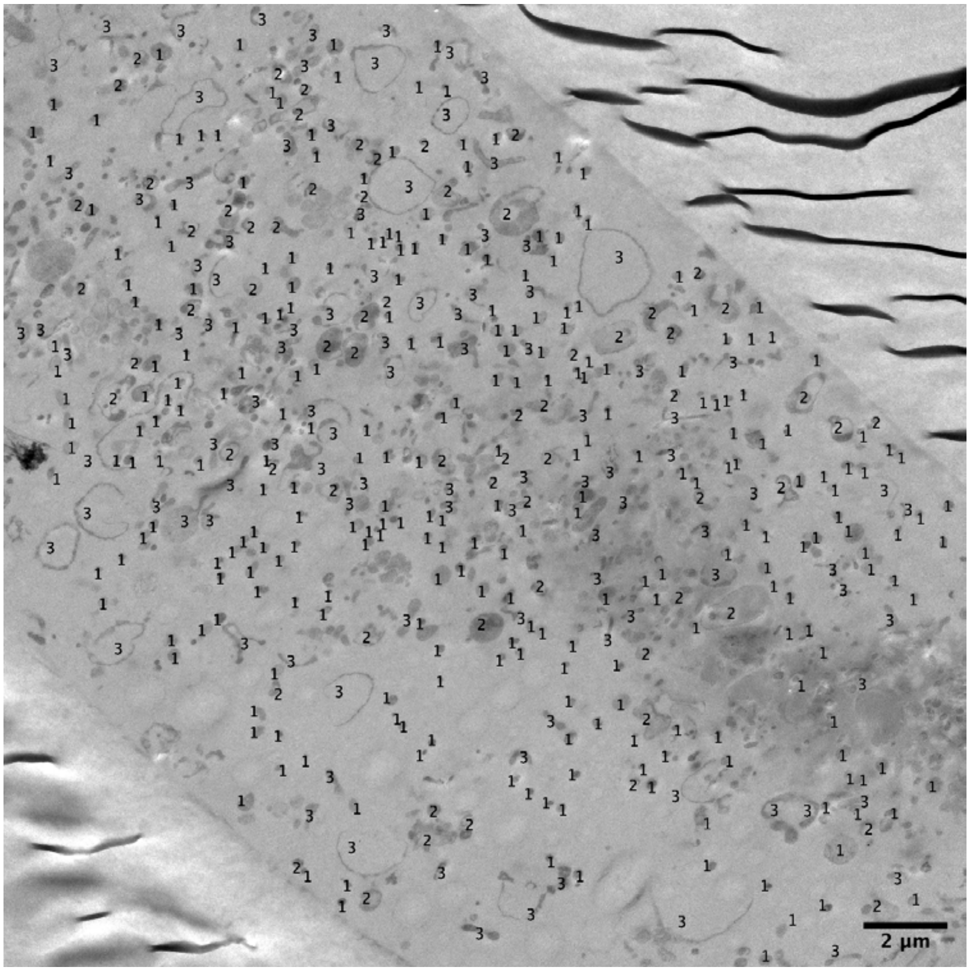

S6b

**POPC overview 1**

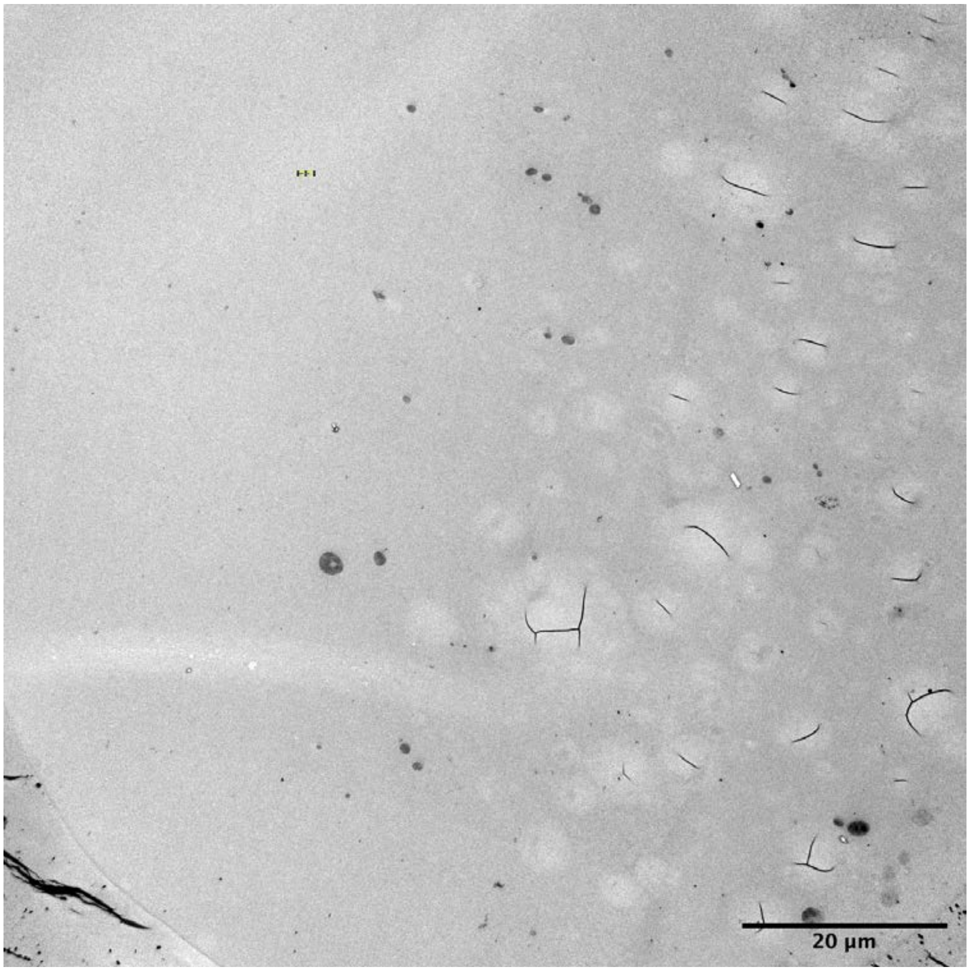


**POPC overview 1 with cell count**

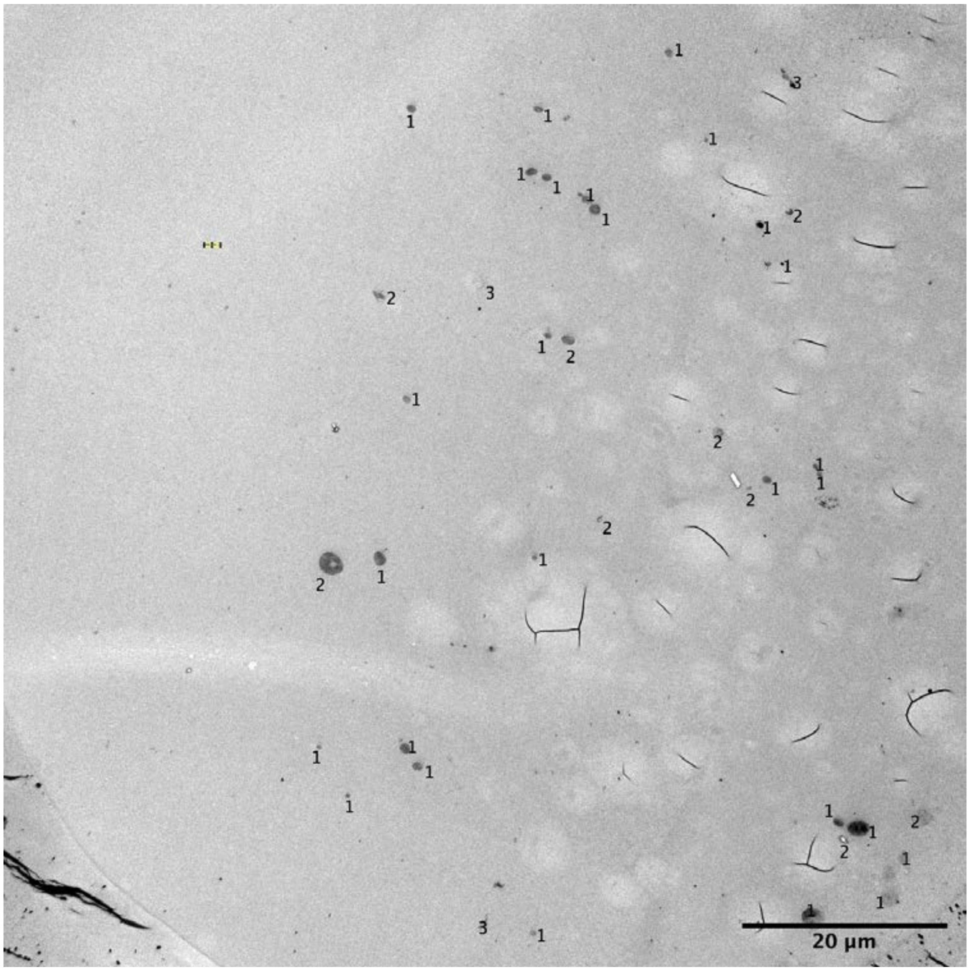

S6c

**POPC overview 2**

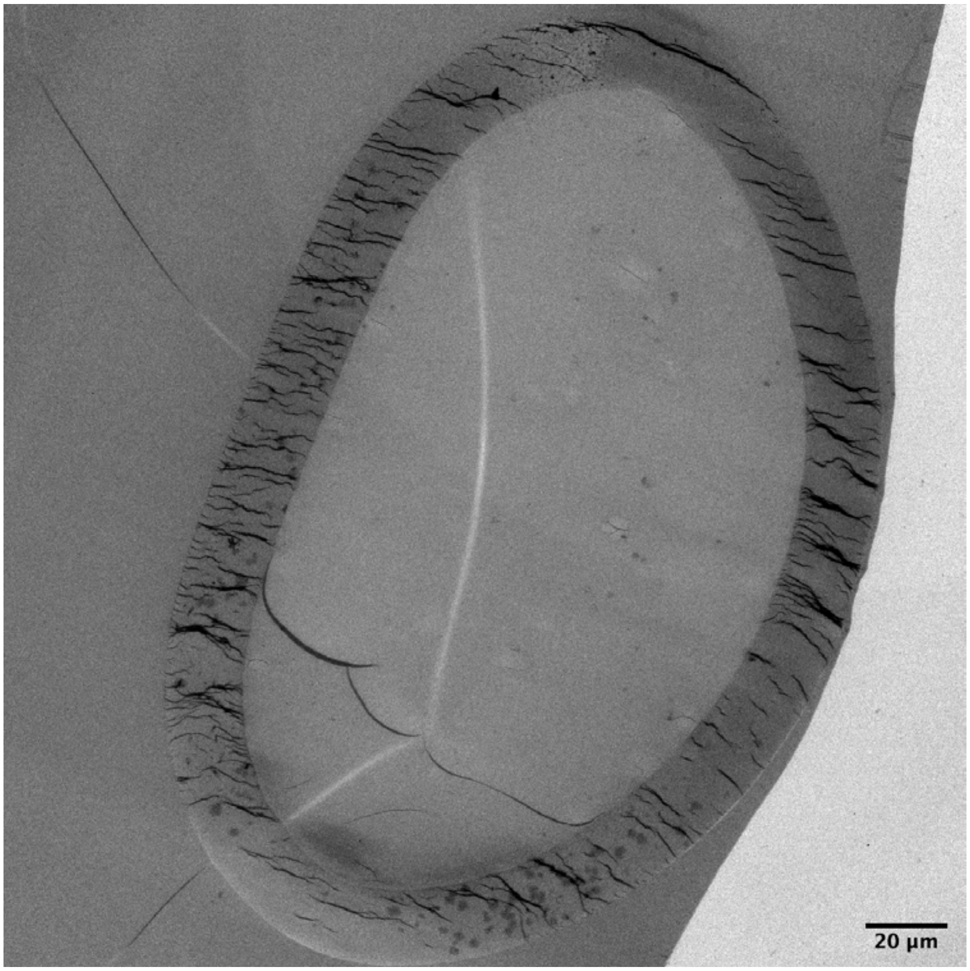


**POPC overview 2
with cell count**

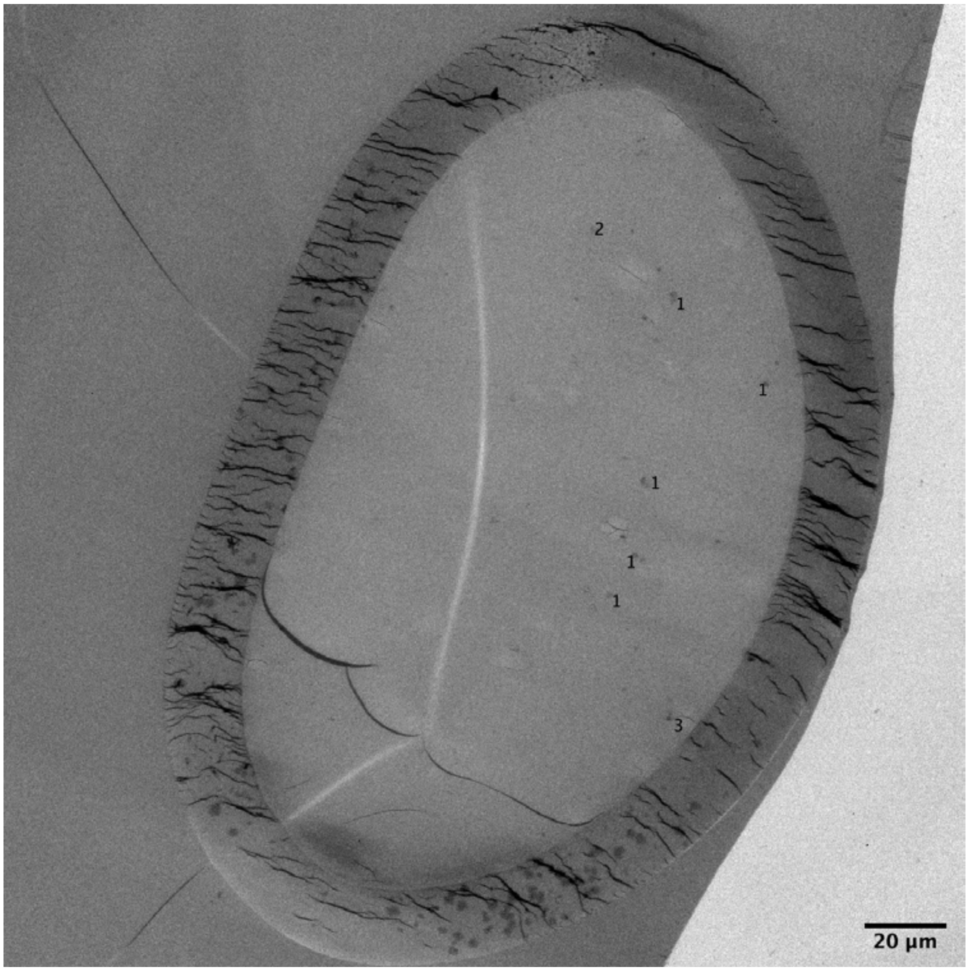

S6d

**POPC overview 3**

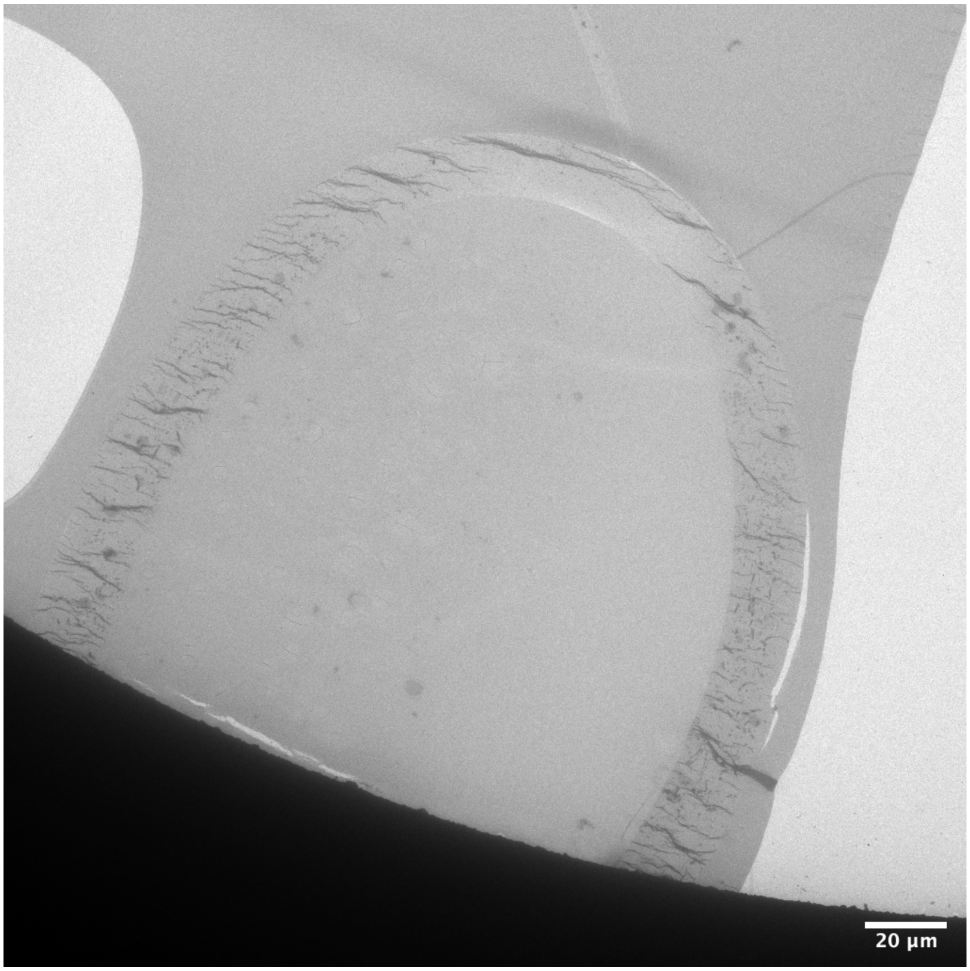


**POPC overview 3 with cell count**

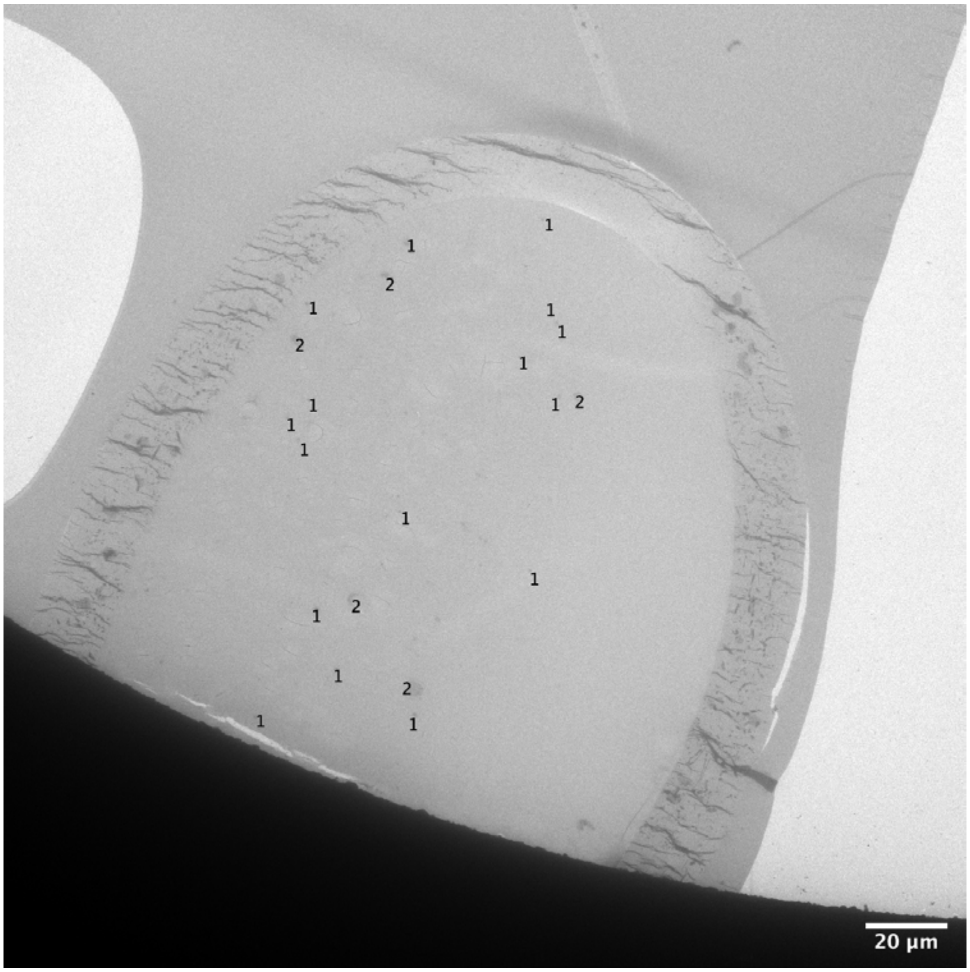

S6e

**D.PC overview 1**

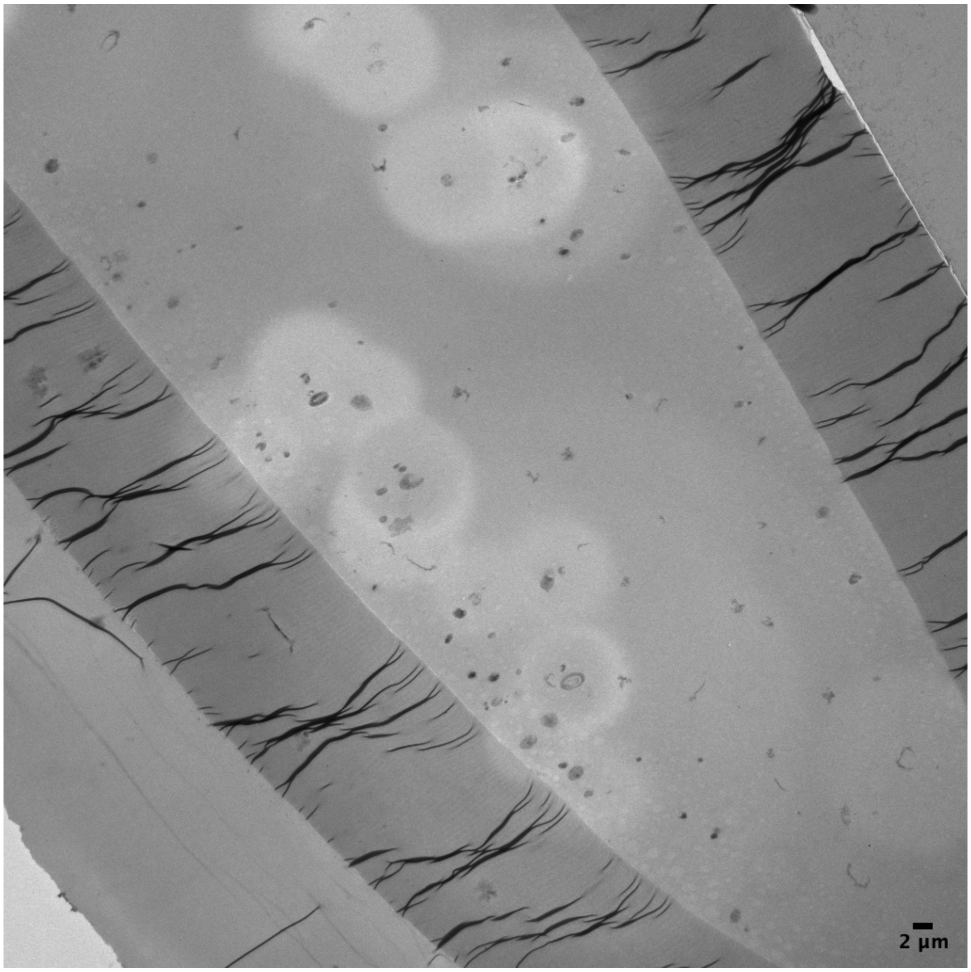


**D.PC overview 1 with cell count**

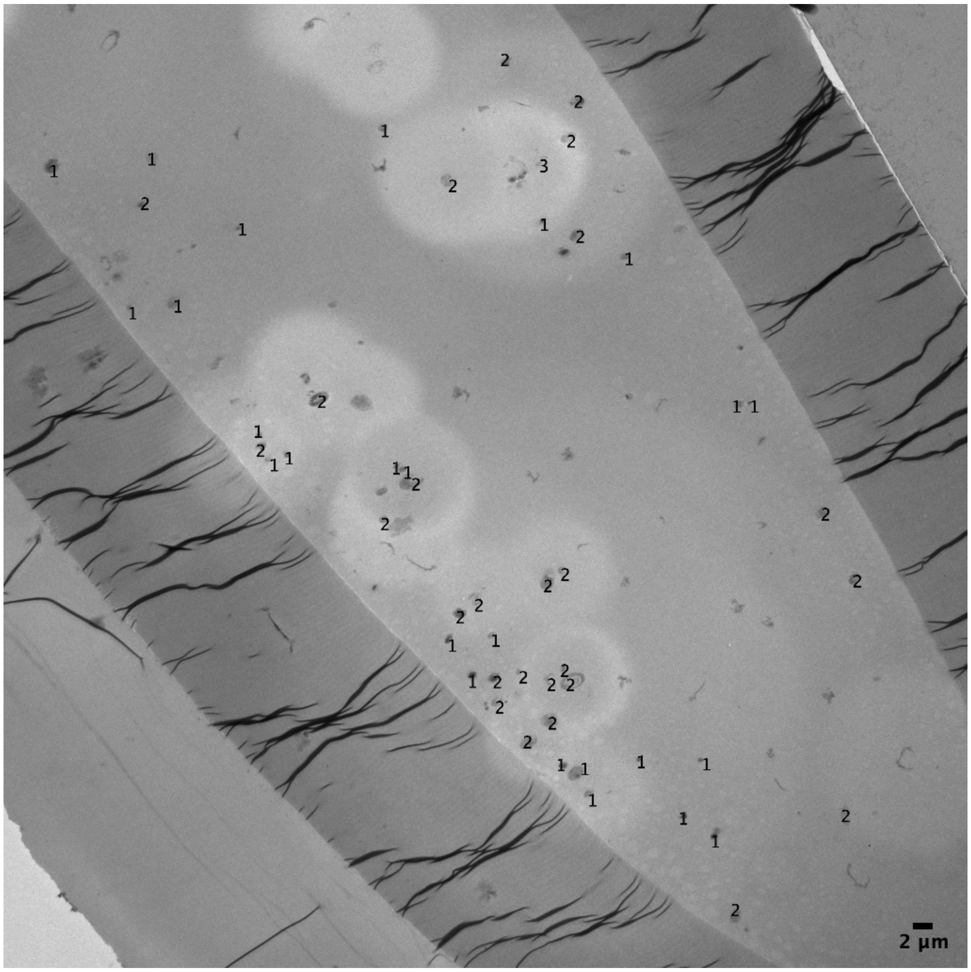

S6f

### D.PC overview 2

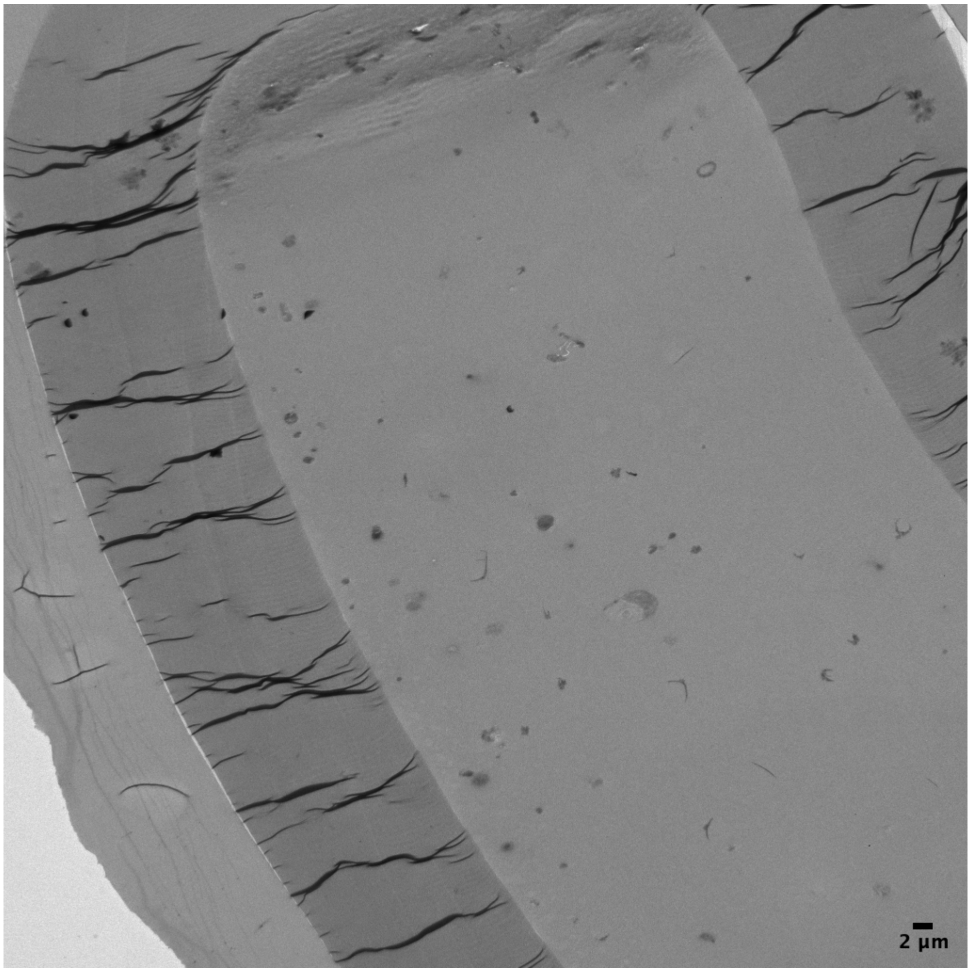


### D.PC overview 2 with cell count

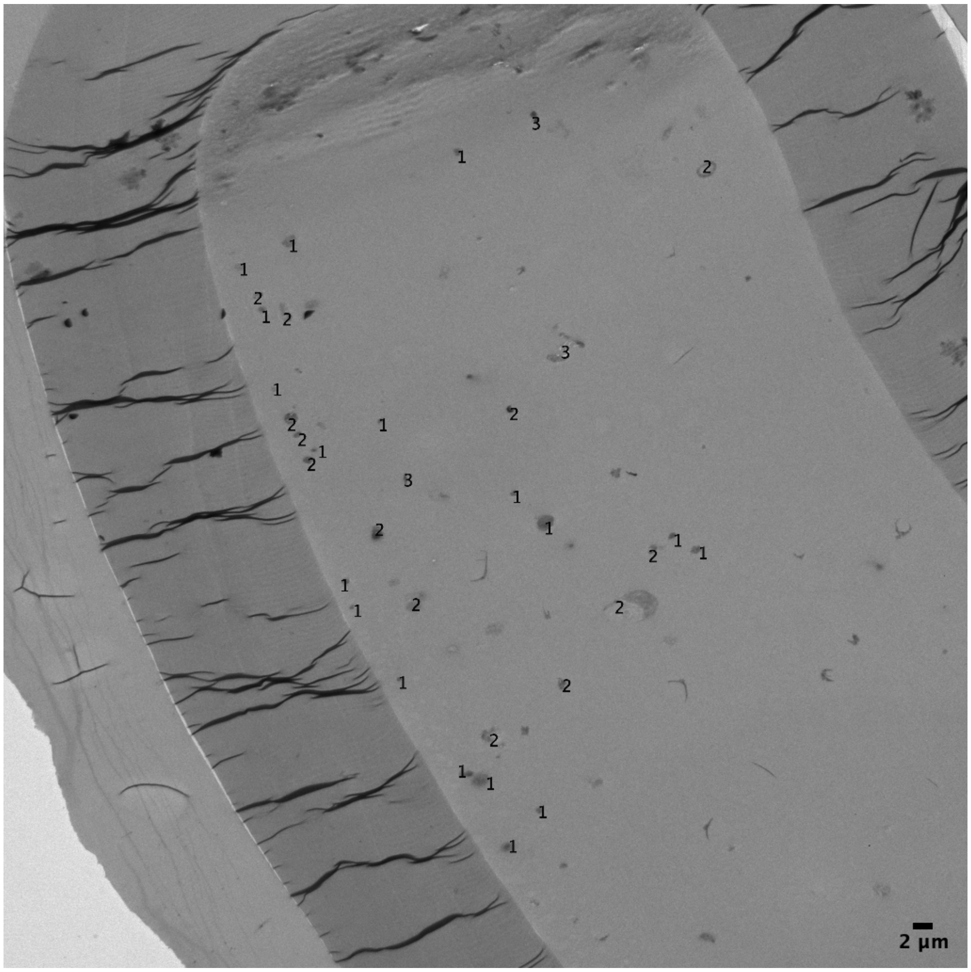

S6g

**D.PC overview 3**

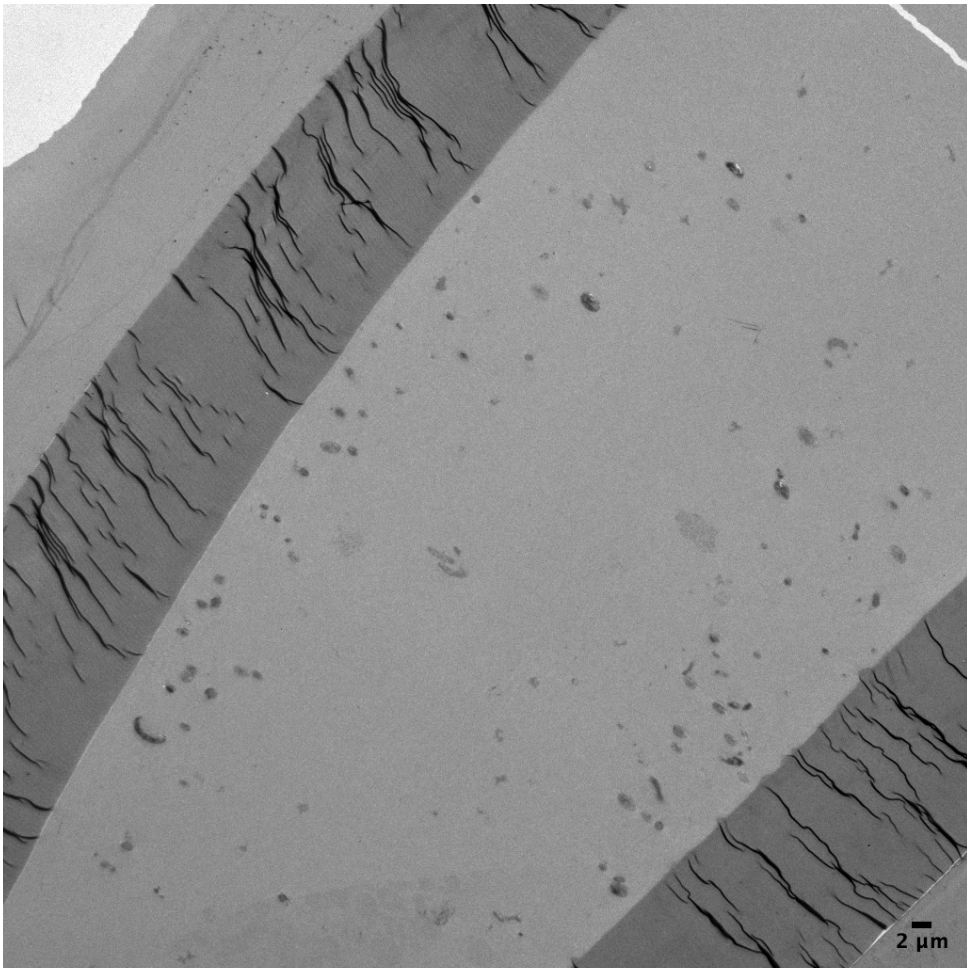


**D.PC overview 3 with cell count**

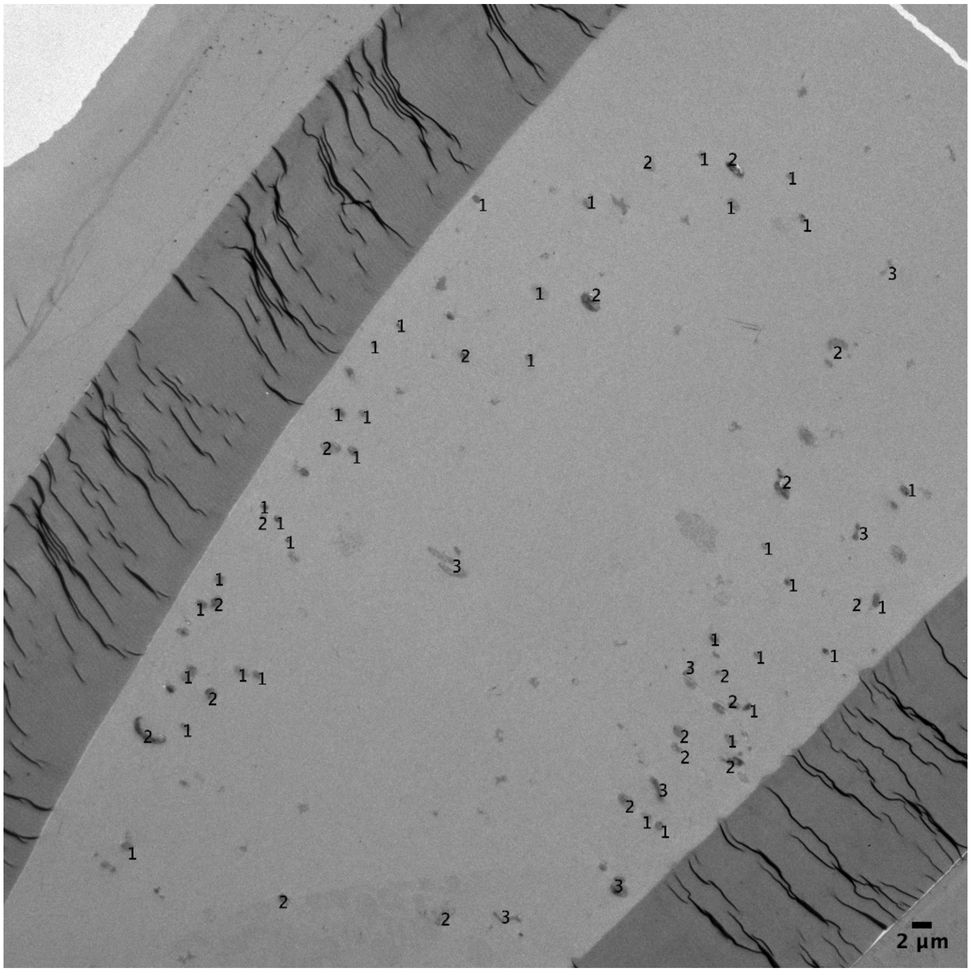

**S6h**

**FBS — Normal** **FBS — Internal Membranes** **FBS — Intracellular Tubules**

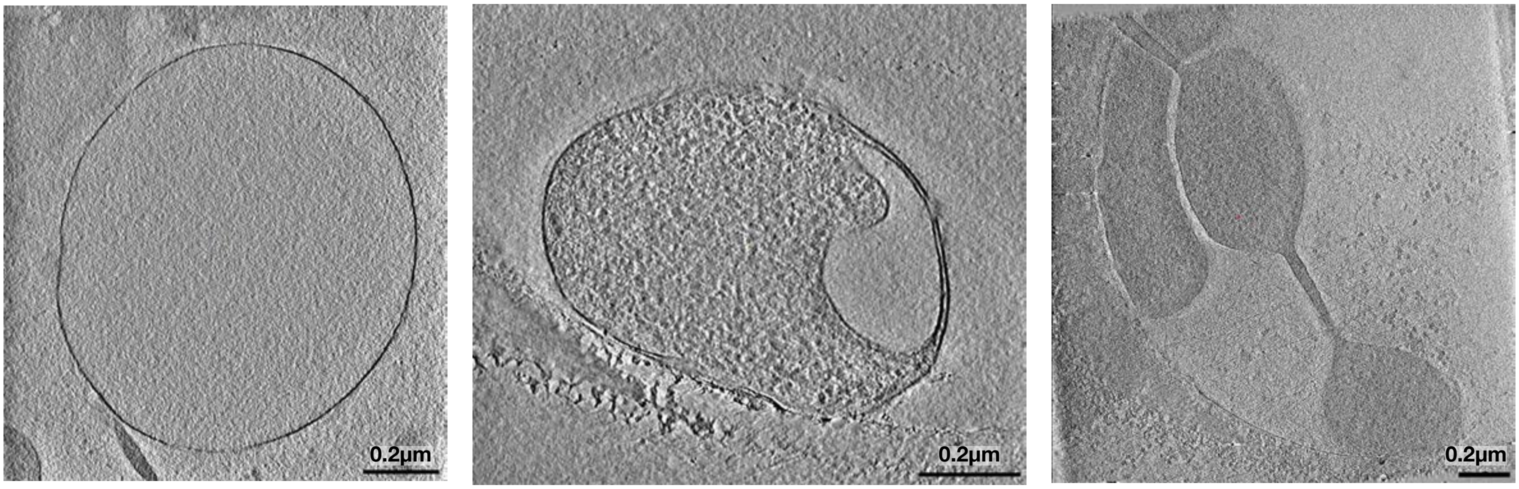


**S6i**

**POPC — Internal Membranes**

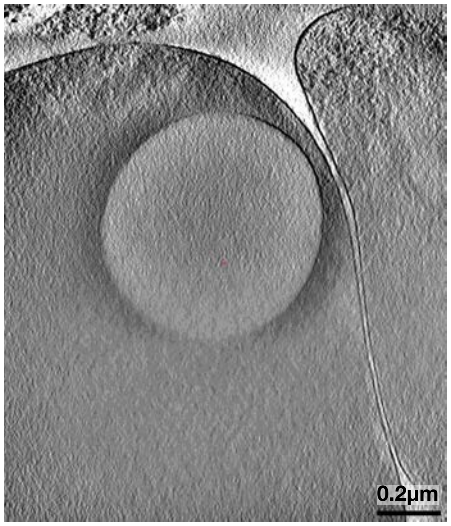

**S6j**

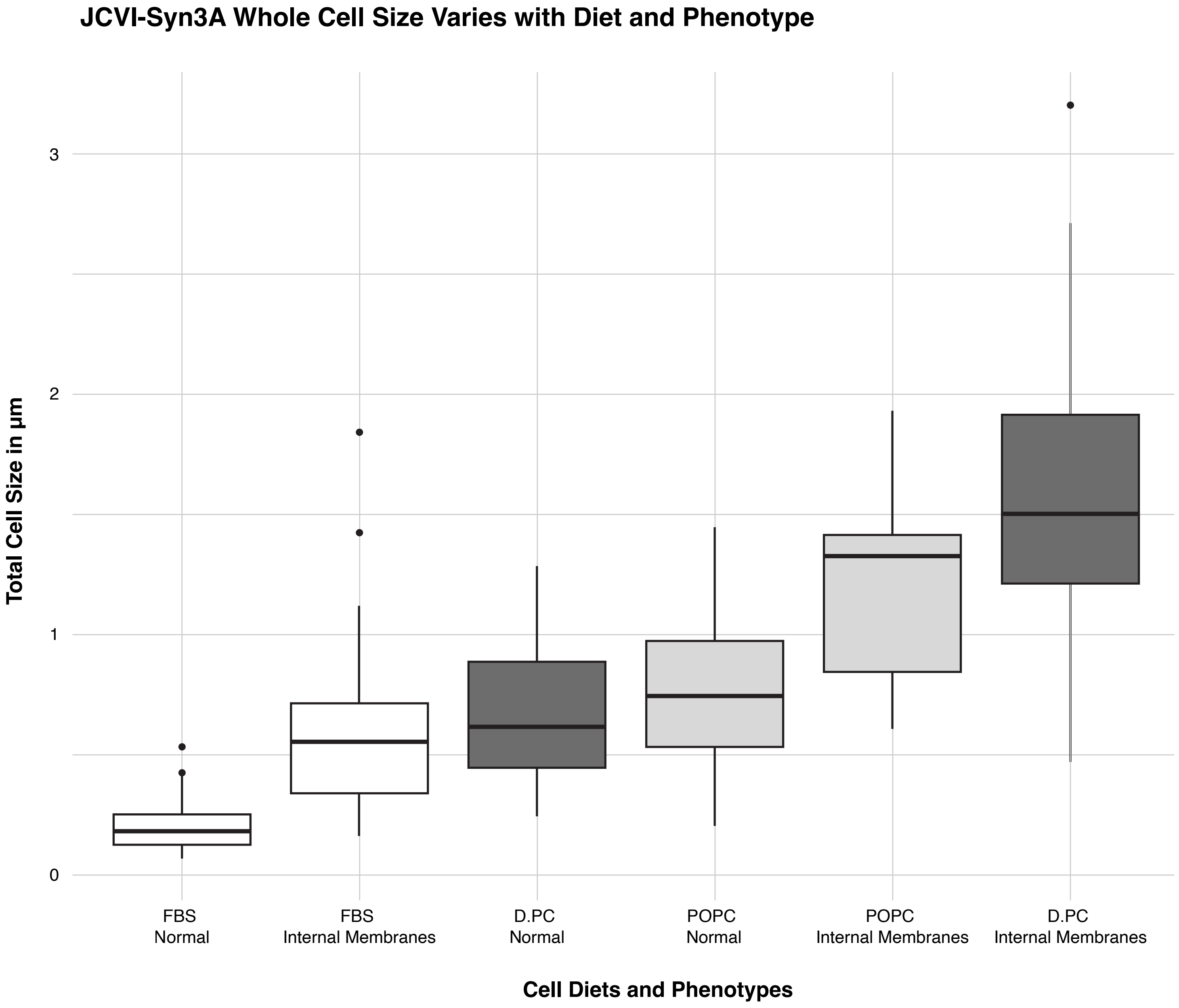


**Figure S6: TEM images of JCVI-Syn3A cells display three distinct phenotypes**

**a-g** TEM overview images of capillaries for JCVI-Syn3A cells on FBS, POPC, and Diether PC diets show a distribution of cells displaying Normal morphology, Internal Membranes, and Intracellular Tubules. For each diet, cells were manually counted to find the number displaying each phenotype. **h** 2D slices of Cryo EM tomograms of FBS cells, displaying cells with the Normal morphology, Internal Membranes, and Intracellular Tubules as well, argue against the possibility that these phenotypes are a fixation artifact. **i** 2D slice of a Cryo EM tomogram of a POPC cell with Internal Membranes. The images of “FBS Internal Membranes” and “POPC – Internal Membranes” in this figure, and the Diether PC cells with Internal Membranes in main Figure 4d, collectively show Cryo EM tomograms of all three diets with internal membranes. **j** Average cell diameters vary as a function of both diet and pheno-type; with Internal Membrane phenotypes displaying both a much larger cell diameter and a larger range of diameters than Normal phenotype cells of the same diet.

**S7a**

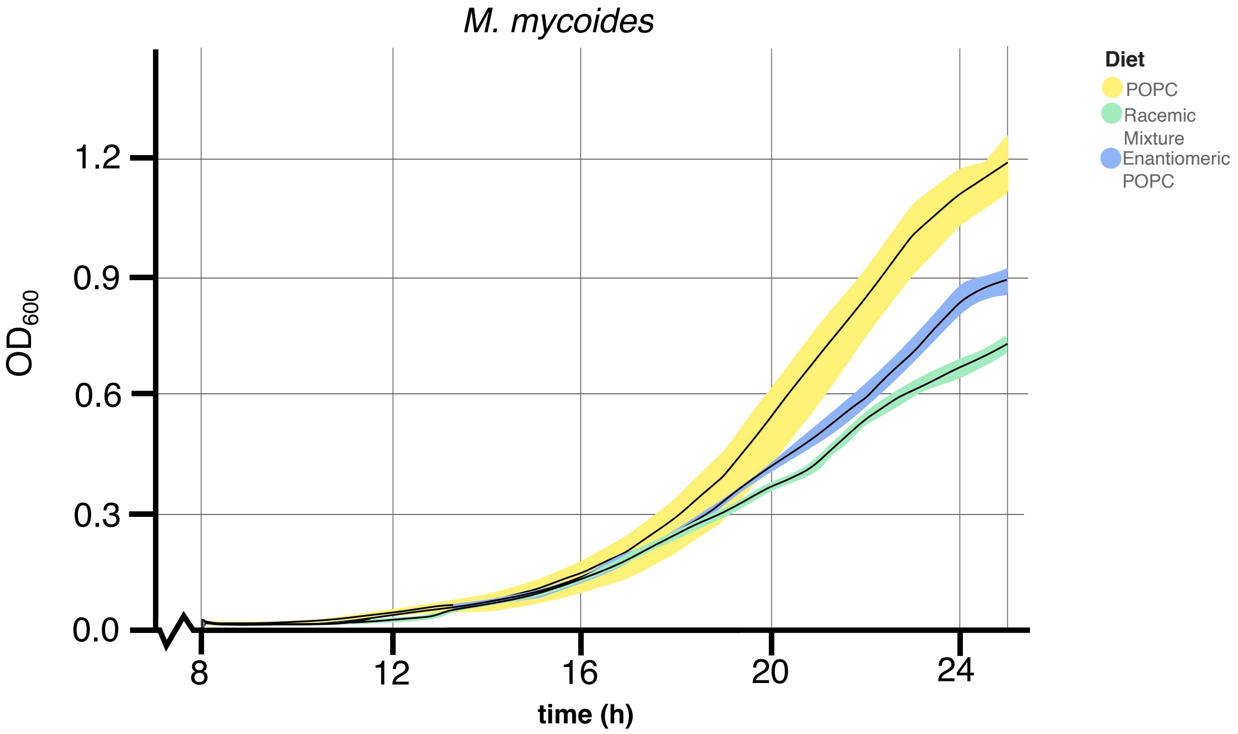


**S7b**

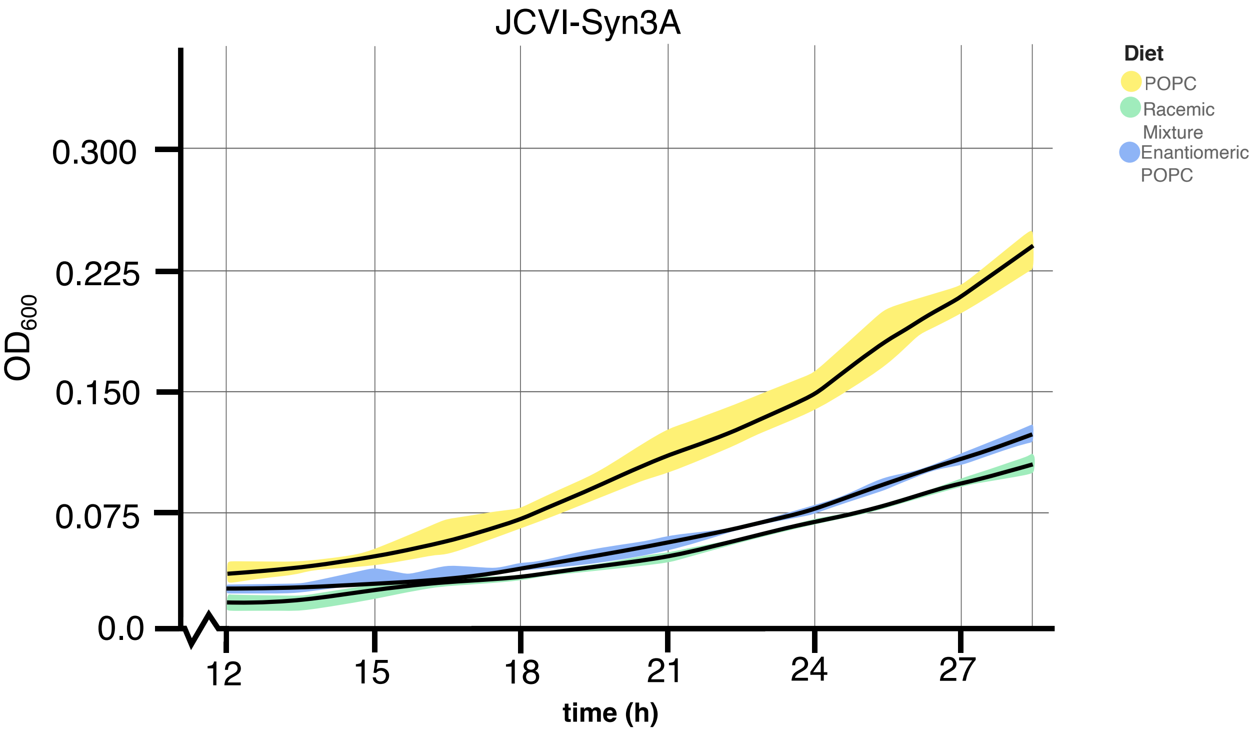


**Figure S7: Growth of *M. mycoides* and JCVI-Syn3A manually measured by $OD_{600}$**

**a** Growth curves for *M. mycoides* on three enantiomeric POPC diets obtained by manual measurements of $OD_{600}$. **b** Growth curves for JCVI-Syn3A on three enantiomeric POPC diets obtained by manual measurements of $OD_{600}$. For both cell types, diets with the enantiomer grow slower than the POPC diet. Complementary phenol red $A_{562}$ growth curves for these diets are shown in Figure S4c.
